# Effects of Parathyroid Hormone, Alendronate and Odanacatib on the mineralisation process in intracortical and endocortical Haversian bone of ovariectomized rabbits

**DOI:** 10.1101/255703

**Authors:** Christina Vrahnas, Pascal R Buenzli, Thomas A Pearson, Brenda L Pennypacker, Mark J Tobin, Keith R Bambery, Le T Duong, Natalie A Sims

**Affiliations:** St. Vincent’s Institute of Medical Research, Fitzroy, Victoria, Australia; The University of Melbourne, Department of Medicine at St. Vincent’s Hospital, Fitzroy, Victoria, Australia; School of Mathematical Sciences, Queensland University of Technology, Brisbane, Australia; MRL, Merck & Co., Inc., West Point, PA, USA.; The Australian Synchrotron, Clayton, Victoria, Australia

**Keywords:** Preclinical Studies, Matrix mineralization, Anabolics, Antiresorptives, Bone histomorphometry

## Abstract

Although cortical bone strength depends on optimal bone composition, the influences of standard therapeutic agents for osteoporosis on bone mineral accrual in cortical bone are not understood. This study compared effects on cortical bone composition of two current therapeutic approaches for osteoporosis: the anti-resorptive bisphosphonate alendronate (ALN), and anabolic intermittent parathyroid hormone (PTH). The experimental anti-resorptive cathepsin K inhibitor, odanacatib (ODN) which inhibits resorption without inhibiting bone formation, was also tested.

To determine effects of these agents on Haversian remodeling and mineral accrual, we compared ALN (100μg/kg/2xweek), PTH(1-34) (15μg/kg, 5x/week) and ODN (7.5μM/day) administered for 10 months commencing 6 months after ovariectomy (OVX) in skeletally mature rabbits by histomorphometry. We used synchrotron-based Fourier-transform infrared microspectroscopy (sFTIRM), coupled to fluorochrome labelling, to measure maturation of the cortical matrix *in situ* at both endocortical and intracortical sites of bone formation.

PTH and ODN, but not ALN, treatment increased bone toughness, and PTH treatment stimulated bone formation, not only on endocortical and periosteal bone, but also in intracortical pores. In Sham and OVX rabbits, normal matrix maturation was observed at both endocortical and intracortical sites including: mineral accrual (increasing mineral:matrix), carbonate substitution (carbonate:mineral) and collagen molecular compaction (amide I:II) *in situ* in endocortical and intracortical bone. ALN treatment reduced bone formation on these surfaces. In ALN-treated bone, while intracortical bone matured normally, endocortical bone did not show a significant increase in mineral:matrix. ODN treatment resulted in slower mineral accrual and limited carbonate substitution. While PTH-treatment did not modify matrix maturation in endocortical bone, the initial stages of mineral accrual were slower in intracortical bone.

In conclusion, these three classes of therapy have differing effects on both bone formation, and the process of bone matrix maturation. ALN suppresses bone formation, and the normal process of matrix maturation in endocortical bone. ODN does not suppress bone formation, but limits mineral accrual. PTH stimulates bone formation, and the matrix formed matures normally in endocortical bone. The ability of PTH treatment to stimulate bone formation in intracortical bone may provide a novel additional mechanism by which PTH increases bone strength.

## Introduction

Postmenopausal osteoporosis is caused by an imbalance between bone formation and resorption such that the level of bone resorption outstrips bone formation, resulting in bone loss and increased fracture risk. Therapies for osteoporosis use two primary approaches to address this imbalance: reducing bone resorption, or promoting bone formation.

Most current long-term therapies for postmenopausal osteoporosis target bone resorption. The most common therapy is bisphosphonate administration which inhibits the activity of bone resorbing osteoclasts [1]. An alternative is denosumab, an antibody to RANKL, which inhibits osteoclast differentiation [2]. These standard osteoclast inhibitors, or anti-resorptives, not only suppress resorption, but also reduce bone remodelling initiation; and due to reduced coupling of bone formation to resorption, anti-resorptives also lower osteoblast-mediated bone formation [3]. Suppressed bone remodelling due to anti-resorptive treatment allows a longer secondary mineralisation period, and bone deposited during treatment continues to mineralize [4]. This has been suggested to result in microdamage accrual, and increased mineral homogeneity in the skeleton [5, 6].

The cathepsin K inhibitor odanacatib (ODN) impairs osteoclastic collagen removal during resorption and has been suggested to provide a therapeutic advantage in bone over standard antiresorptives, such as bisphosphonates. This has been suggested because bone formation was not inhibited by ODN during remodeling in preclinical and clinical biopsy studies, and was even increased in the modelling periosteum in ovariectomized monkeys [7, 8] and transiliac biopsies from ODN-treated patients [9]. There is little data to indicate how cathepsin K inhibition affects bone matrix composition. Lifelong cathepsin K ablation in a genetically altered mouse was associated with altered bone composition: hypermineralization (greater mineral:matrix ratio) in midshaft cortical bone, but not in trabecular bone[10]. This was presumed to occur due to prolonged secondary mineralisation, but initial (primary) mineralisation may also be more rapid when cathepsin K activity is impaired. These findings in the murine genetic model had four major limitations: (a) cathepsin K inhibition was in effect throughout skeletal growth and development, (b) murine bone is non-Haversian, (c) the mice were young and still rapidly growing, and (d) the method used to measure bone composition could not distinguish regional changes in composition.

In contrast to these antiresorptives, anabolic therapy, such as intermittent parathyroid hormone (PTH) promotes bone formation, and can therefore increase bone mass [11]. A similar anabolic mechanism has been suggested for abaloparatide, which is a modified N-terminal preparation of PTH-related protein (PTHrP)[12], recently FDA-approved for osteoporosis therapy. PTH increases bone mass both by increasing bone formation on active bone remodelling surfaces that have undergone prior resorption, and by stimulating new bone formation on quiescent bone surfaces, sometimes termed modelling surfaces [13]. This has been noted both on trabecular and endocortical surfaces [13, 14]. However, on intracortical surfaces, PTH has been reported to increase porosity by stimulating bone remodeling [15] and the initial remodelling phase (osteoclast recruitment to the bone surface). No analysis of PTH effects on bone formation on modelling intracortical surfaces (lacking prior resorption) has been reported. Although there were early concerns that bone deposited during PTH treatment had lower mineral content than control bone, we recently reported, using a murine model, that the low average mineral content of PTH-treated bone results from a higher proportion of new bone, rather than any change in the quality of bone deposited during PTH treatment on the murine periosteum (outer cortical surface) [16]. This was achieved using a combination of fluorochrome labelling and synchrotron-based Fourier-transform infrared microspectroscopy (FTIRM). By measuring matrix composition at increasing depth on a bone forming surface without recent prior resorption (the murine periosteum), this method also provided a way of identifying and measuring three related processes that occur as the bone matrix matures: (1) mineral accumulates within the matrix (mineral:matrix ratio), (2) carbonate substitution within the mineralised matrix (carbonate:mineral ratio) increases, and (3) collagen molecules become more compact in their perpendicular axis (amide I:II ratio) as mineral accumulates [16].

To determine how these agents influence bone matrix accrual in Haversian cortical bone, we have carried out a side-by-side comparison of three therapies used for postmenopausal osteoporosis with distinct mechanisms (PTH, ALN, and ODN) on bone material composition in ovariectomized rabbits since, unlike smaller mammals, they exhibit Haversian bone remodeling. Because the effects of these agents on the process of bone matrix maturation are not known, we assessed both endocortical and intracortical bone, and used fluorochrome labelling to distinguish regions of persistent modelling-based bone formation (>4 months) from regions containing remodelling‐ and modelling-based bone formation <4 months. We identify that ALN, ODN, and PTH treatments each modify the process of bone matrix mineral accrual in unique ways, and that PTH treatment stimulates bone formation on modelling surfaces of intracortical bone, revealing a novel additional mechanism by which PTH may increase bone strength.

## Materials and Methods

### Animals and treatments

This study was conducted in accordance with recommendations of the Guide for the Care and Use of Laboratory Animals and approved by the Institutional Animal Care and Use Committee of MRL, West Point. NZW rabbits were obtained from Covance, Inc. (Denver, PA) and singly-housed in wire-bottomed cages under standard laboratory conditions with a 12-12 hour light cycle. Rabbits were administered daily 125 gm of Harlan Teklad 2030 Global rabbit diet (Harlan Laboratories, Inc., Madison, WI) and water *ad libitum*. Rabbits were subjected to either sham or ovariectomy (OVX) surgery at 7 months of age. At 6 months post-surgery, OVX rabbits were randomized by lumbar vertebrae 5-6 bone mineral density and body weight and treated for 10 months with alendronate (ALN, Merck, 100 μg/kg/2xweek, by subcutaneous injection), odanacatib (ODN, Merck, 7.5 μM•24 hr/day in Harlan Teklad 2030 and 2031 control diet) or human parathyroid hormone (1-34) (PTH, Bachem, 15 μg/kg/day, s.c. 5 times per week) **(Figure 1A)**. Due to palatability issues, all rabbits were switched to global rabbit diet 2031 (Harlan Teklad) after six months of treatment. After 6-months into the treatment period, two injections of oxytetracycline (T, Oxyvet 100 LP, Vetoquinol, Canada, 30 mg/kg IM) were administered with a seven-day interval. After 10-months of treatment, a second double-labeling with calcein (Sigma, 8 mg/kg) was also given on the tenth (C1) and third (C2) day before necropsy **(Figure 1A)**. Age-matched sham operated rabbits were also used as controls.

**Figure 1:**
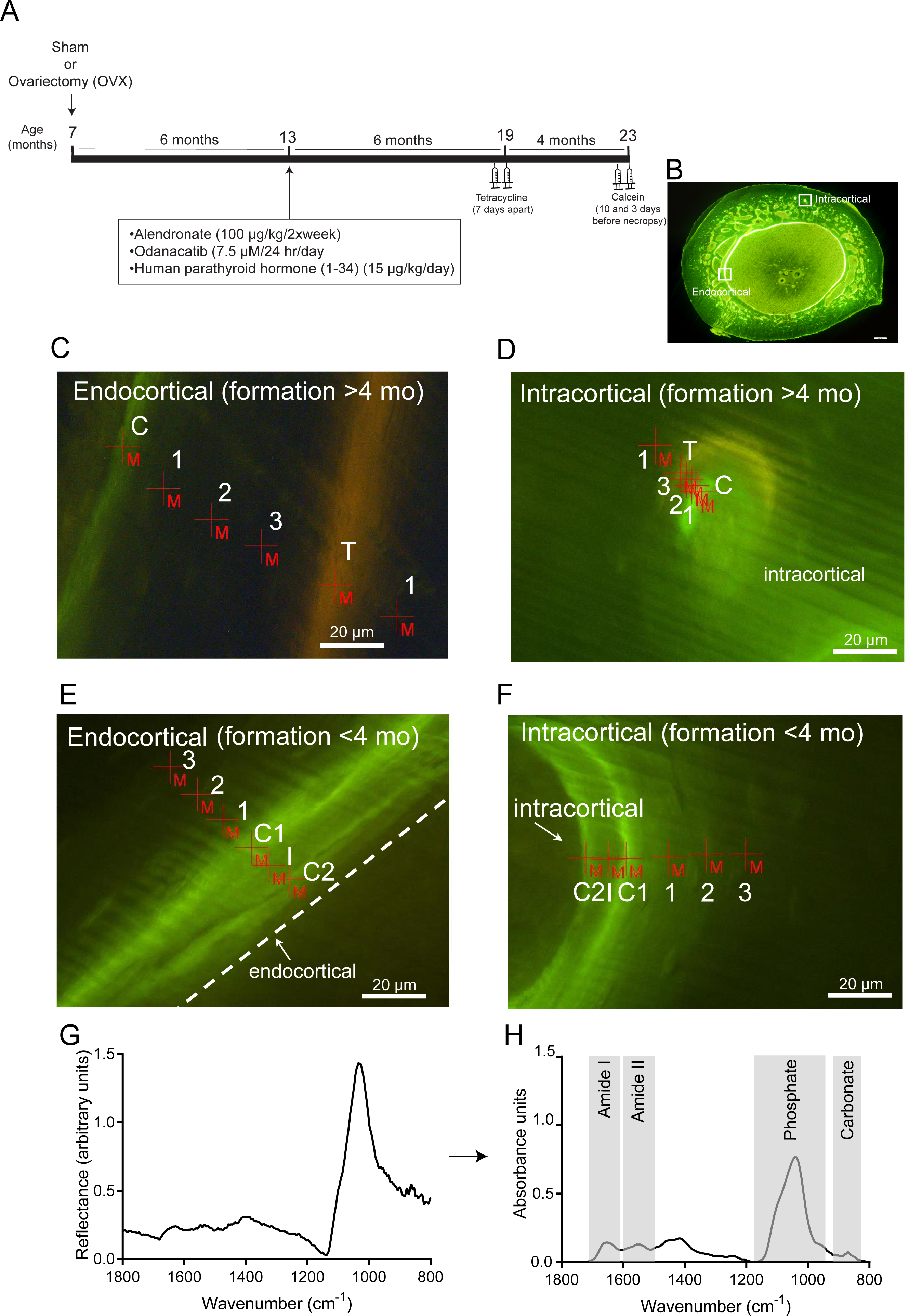
A: Schematic showing treatment protocol for this study. B-F: Representative fluorescent images from PTH-treated samples showing overall image (B) and regions of measurement (red +M), and fluorochrome labels at endocortical (C,E) and intracortical (D,F) sites with parallel tetracycline and calcein label or lacking underlying calcein label. Labels are: dual calcein labels (C1, C2), intermediate region between calcein labels (I), tetracycline label (T), and regions measured at increasing depth into the cortex (numbers 1-3). Scale bar = 20 microns. G,H: Representative images showing (G) raw reflectance spectrum before mathematical processing, (H) the same spectrum after processing, and approximate regions selected for analysis of amide I, amide II, carbonate and phosphate.

### Bone mineral density and histomorphometry

For all endpoint measurements, the investigators were blinded during analysis. For *ex-vivo* densitometric analysis, the 3^rd^ and 4^th^ lumbar vertebrae were fixed in 70% ethanol and scanned by dual-energy X-ray absorptiometry (DXA) using small animal acquisition high resolution software on an Hologic Discovery A fan-beam bone densitometer (Hologic, Inc., Waltham, MA). Regions of interest were drawn around the vertebral body, excluding transverse processes. Bones were immersed in an acrylic box in 50 mm of water. The vertebrae were divided into two regions: region 1 (whole vertebra) and region 2 (a box placed in the middle 35% of the vertebra, with 30% remaining at the cranial end, and 35% remaining at the caudal end). Region 2 was excluded from analysis since this region routinely does not contain any trabecular bone; data presented therefore represent cortical BMD.

For Fourier-Tranform Infrared Microspectroscopy (FTIRM) analysis and histomorphometry, right femora were collected and fixed in 70% ethanol. Femoral length was measured to obtain the midpoint. A cortical section extending 0.5 cm both proximal and distal to the midpoint was removed using an Exakt 300 CP bandsaw (Exakt Technologies, Inc., Oklahoma City, OK) and embedded undecalcified in 90% methyl methacrylate/10% dibutyl phthalate. Sections from the distal end of the cross-section were cut approximately 100μm thick using a Leica 1600 microtome (Leica Instruments GmbH; Nussloch, Germany). Cross-sections for FTIRM were stored between 2 Whatman filter papers and glass slides before shipping and analysis. Sections for histomorphometry were mounted onto glass slides using Eukitt’s mounting media.

For histomorphometry, a light/epifluorescent microscope (Optiphot II, Nikon, Japan) equipped with an Optronics DEI-750 CE (Tuttlingen, Germany) video camera interfaced to Bioquant Osteo Software (Bioquant Image Analysis Corp., Nashville, TN), was used to collect the raw data. Standard nomenclature for bone histomorphometry was used [17]. Histomorphometric endpoints representing endocortical mineralizing surface (Ec.MS/BS, %), mineral apposition rate (Ec.MAR, μm/d), and bone formation rate (Ec.BFR/BS, μm^3^/μm^2^/yr) and intracortical Haversian mineralizing surface (H.MS/BS, %), mineral apposition rate (H.MAR, μm/d), and bone formation rate (H.BFR/BS, μm^3^/μm^2^/yr) were calculated. Mineralizing surface was calculated based on double-labeled surface plus half single-labeled surface divided by total bone surface. Ec.MAR and H.MAR were calculated from double-labeled width and interlabel time period. Periosteal mineralizing surface (Ps.MS/BS, %), mineral apposition rate (Ps.MAR, μm/d), and bone formation rate (Ps.BFR/BS, μm^3^/μm^2^/yr) were calculated for both interim tetracycline labels and end of study calcein labels. Bone surfaces with insufficient double label were excluded from the calculation of mean mineral apposition rate (Ec.MAR, H.MAR and Ps.MAR). Because most samples exhibited double label on the endocortical surface, and since reference values have not yet been developed for rodent MAR in low turnover situations, we used two approaches to calculate Ec.BFR/BS (reported separately in Figure 2 and Table 1): (i) according to ASBMR recommendations [18] animals with no double label were assigned a bone formation rate of zero and in those samples where double calcein labels were detected but were insufficient to measure endocortical mineral apposition rate (Ec.MAR), values of 0.1μm/day were imputed and used for Ec.BFR/BS calculations, (shown in Figure 2), and (ii) samples without double label were recorded as “missing data” and Ec.MAR and Ec.BFR/BS calculated with those samples excluded (shown in Table 1) as recommended by Recker et al [19].

**Figure 2:**
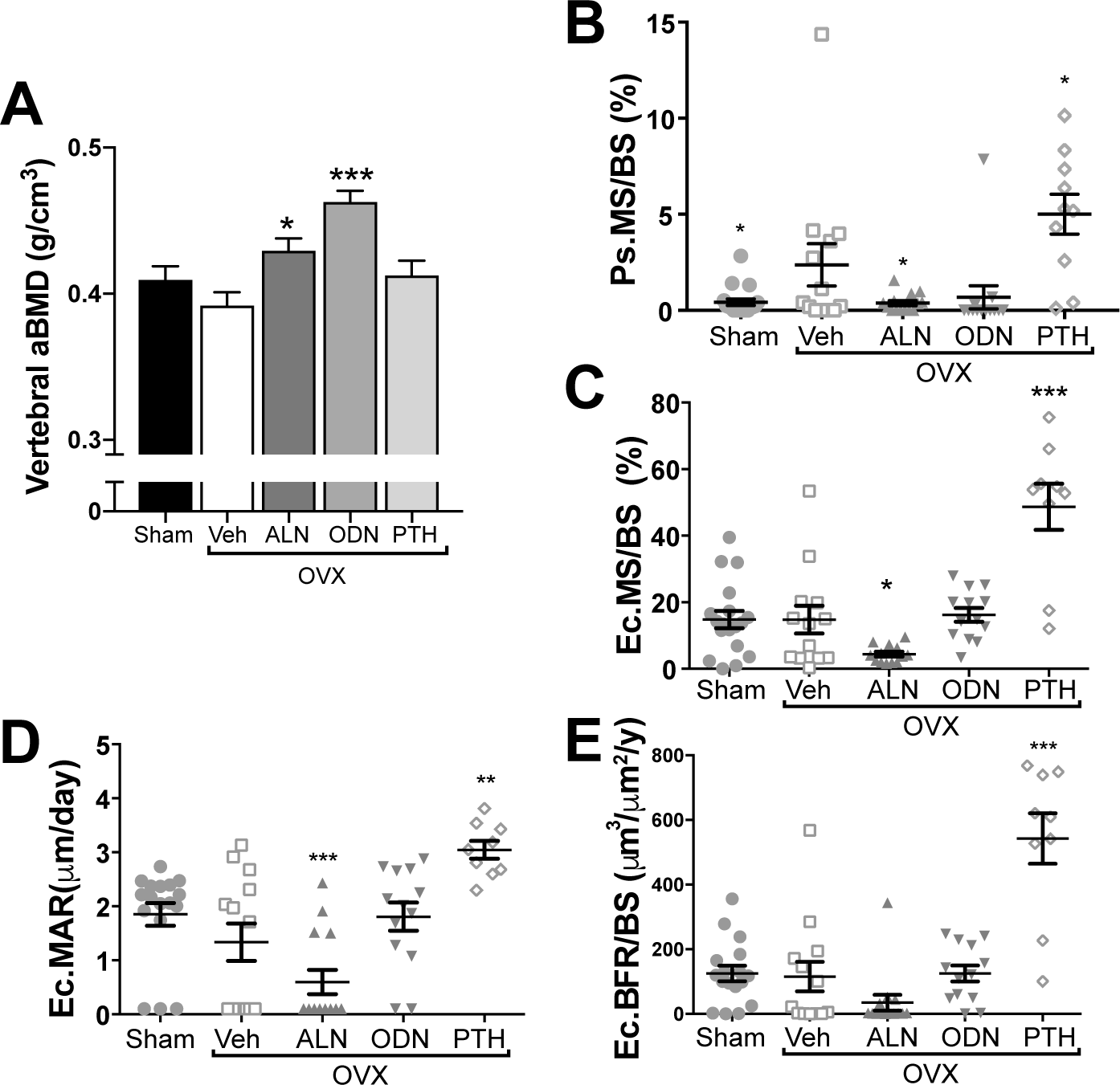
Lumbar vertebral bone mineral density (BMD) (A) and histomorphometric analysis of the femoral diaphyseal periosteum (B) and endocortical surface (C-E) from sham-operated and ovariectomized (OVX) rabbits treated with vehicle, alendronate (ALN), odanacatib (ODN) or parathyroid hormone (PTH). PsMS/BS = periosteal mineralising surface, EcMS/BS = endocortical mineralising surface, EcMAR = endocortical mineral appositional rate (with imputation), EcBFR/BS = endocortical bone formation rate (with imputation). Data is mean ± SEM; n= 10-17 per group. *, p<0.05, **p<0.01 vs OVX+vehicle.

**Table 1:** Biomechanical strength testing and dynamic histomorphometric analysis of intracortical Haversian bone, and of the central femoral cortex. H=Haversian; MS/BS = mineralising surface / bone surface; MAR = mineral appositional rate. BFR/BS = bone formation rate / bone surface. Values are mean  SEM; ^+^, p<0.05 vs Sham; **, p<0.001, **, p<0.01; *, p<0.05 vs OVX-vehicle; H.MAR could not be analysed statistically due to insufficient label in most samples.

|  | Sham | OVX+Vehicle | OVX+ALN | OVX+ODN | OVX+PTH |
| --- | --- | --- | --- | --- | --- |
| **N - biomechanical testing** | 19 | 13 | 14 | 13 | 12 |
| *Extrinsic* |  |  |  |  |  |
| Peak Load (N) | 364 ± 11 | 422 ± 22<sup>+</sup> | 388 ± 19 | 452 ± 17 | 477 ± 23 |
| Stiffness (N/mm) | 617 ± 80 | 691 ± 49 | 660 ± 27 | 673 ± 27 | 684 ± 47 |
| Work-to-Failure (N-mm) | 219.7 ± 14.7 | 229.3 ± 21.1 | 220.3 ± 23.2 | 304.2 ± 17.9* | 318.7 ± 26.5** |
| *Intrinsic* |  |  |  |  |  |
| Ultimate Stress (MPa) | 149.0 ± 5.3 | 156.3 ± 4.7 | 150.4 ± 6.9 | 163.7 ± 6.5 | 160.2 ± 4.7 |
| Modulus (MPa) | 12321 ± 1624 | 16664 ± 5350 | 12081 ± 466 | 11267 ± 422 | 10630 ± 776 |
| Toughness (MPa) | 1.89 ± 0.13 | 1.85 ± 0.14 | 1.80 ± 0.18 | 2.37 ± 0.14* | 2.34 ± 0.18* |
| **N - histomorphometry** | 18 | 13 | 14 | 13 | 11 |
| N with Ec. double labels | 14 | 7 7 | 4 4 11 | 11 11 | 11 |
| Ec.MAR ($\mu\text{m}/\text{d}$ without imputation) | 2.23 ± 0.07 | 2.39 ± 0.20 | 1.84 ± 0.22 | 2.12 ± 0.18 | 3.00 ± 0.14* |
| Ec.BFR/BS ($\mu\text{m}^3/\mu\text{m}^2/\text{yr}$ without imputation) | 118.0 ± 23.6 | 114.5 ± 46.2 | 33.9 ± 24.4 | 124.8 ± 24.8 | 544.8 ± 64.1*** |
| N with sufficient H. (intracortical) double labels | 1 | 2 | 1 | 1 | 9 |
| H.MS/BS (%) | 0.35 | 0.50 ± 0.22 | 0.84 | 1.48 | 11.46 ± 2.61*** |
| H.MAR ($\mu\text{m}/\text{d}$) | 3.04 | 2.53 ± 0.34 | 2.13 | 2.13 | 2.65 ± 0.10 |
| H.BFR/BS ($\mu\text{m}^3/\mu\text{m}^2/\text{yr}$) | 0.9 | 2.7 ± 1.8 | 2.5 | 2.7 | 104.4 ± 28.4*** |

### Bone Strength Testing

Left femora were cleaned of soft tissue and were retained frozen (−20°C) until testing. Bones were thawed overnight at 4°C before being tested. Biomechanical evaluation was performed using the 858 Mini Bionix Servohydraulic Test System, Model 242.03. All data were collected using TestWorks^®^ version 3.8A for TestStar^TM^ software, version 4.0C. The femurs were tested in three-point bending at 1mm/second. The whole femur was positioned with the anterior side up on stationary lower supports. Peak load, stiffness and area under the curve (AUC) were determined from the load-displacement curve. Ultimate stress was calculated using the maximum load, span, perimeter (to calculate the radius) and cross-sectional moment of inertia (CSMI), as determined by pQCT. Modulus was calculated using the stiffness, CSMI and the span. Toughness was calculated using the AUC, specimen diameter, CSMI and the span. After testing, the broken femur was examined to ascertain fracture direction and location.

### Synchrotron-based Fourier-transform infrared microspectroscopy (sFTIRM)

sFTIRM data were collected using a Bruker Hyperion 2000 IR microscope coupled to a V80v FTIR spectrometer (Bruker Optik, Ettlingen Germany) located at the IR Microspectroscopy beamline at the Australian Synchrotron. The Hyperion microscope was equipped with an epifluorescence accessory suitable for calcein visualisation (excitation/emission ~490nm/ ~515 nm).

100 μm sections were placed on a 45 mm diameter × 3 mm cesium iodide (CsI) disks (Crystan Limited, Poole, UK) for spectral imaging. Adjacent regions were imaged in endocortical and intracortical bone (**Figure 1B**) using reflection-based sFTIRM[20], with a clear region of the CsI window used to acquire a reflectance reference spectrum. Regions containing fluorochrome labels were selected to allow us to define “bone-age”-matched spectral measurements. 9 randomly selected samples per treatment group were scanned in multiple adjacent 20μm x 20μm measurement regions at fluorescent label sites and at increasing depth into the older bone (**Figure 1C-F**). Spectra were collected from two types of bone forming surfaces: (1) regions of calcein labels with underlying parallel tetracycline label and therefore lacking recent prior resorption (formation >4 months; modelling bone) (**Figure 1C,D**), and (2) regions with calcein labels on a surface lacking underlying tetracycline label (bone formation < 4 months that is a mix of recent modelling-based bone formation and remodelling-based bone formation, where the tetracycline label was resorbed) (**Figure 1E,F)**. Since measurements and categorization were performed on uncoverslipped sections to allow FTIR microanalysis, it was not possible to distinguish between modelling and remodelling bone by assessing whether the underlying cement line was scalloped or smooth. In regions with underlying tetracycline label, spectra between the calcein and tetracycline labels were spaced evenly to match bone age in the samples despite different apposition rates. Where there was no underlying tetracycline label, spacing was matched to the distance between the calcein labels. Since periosteal surfaces demonstrated little mineralising surface, composition of newly formed bone on the periosteum was not measured. Although many sections from non-PTH-treated bones lacked sufficient intracortical double label for histomorphometric analysis, we were able to identify calcein labelled regions in most sections prepared for spectroscopic analysis by sFTIRM (see Table 2). Spectra were collected in specular reflection geometry from the specimen surface, over the 0 cm^−1^−3800 cm^−1^ spectral range using a mercury cadmium telluride detector, at 4 cm^−1^ spectral resolution and 512 co-added scans/pixel spectral resolution. All data acquisition was undertaken with OPUS version 6.5 and data analysis completed with OPUS version 7.2 (Bruker Optik, Ettlingen Germany). After acquisition, Kramers-Kronig transformation (KKT)[20] was applied to reflectance data (**Figure 1F,G**); this enables retrieval of the phase shift during reflection, from which the sample’s absorption properties can be deduced. Phase retrieval by KKT is a sensitive operation requiring mathematical integration of the reflectance spectrum over all frequencies from zero to infinity. We used an in-house numerical procedure implemented in MATLAB to extrapolate reflectance data outside the experimental range and to perform the KKT[21]. Each reflectance spectrum was first normalised to a 0-1 amplitude by correcting for CsI background, and smoothed by a 11-point moving mean, corresponding to a 40 cm^−1^ averaging window. These data were extended at far infrared (IR) frequencies <960 cm^−1^ by a template bone IR spectrum obtained from a polished mouse bone block in a prior publication (Figure 3A in [20]). It was digitised and merged with the reflectance data after scaling and shifting so that 1) the phosphate peak height ratios at 557 cm^−1^ and at 1025 cm^−1^ in the template IR spectrum were maintained in the merged spectrum; and 2) the phosphate minimum height at 380 cm^−1^ in the template IR spectrum matched the reflectance data baseline in the 1500 cm^−1^–2000 cm^−1^ region. The transition at 960 cm^−1^ was smoothed by linear superposition of the reflectance data and of the scaled template IR spectrum in the range 920 cm^−1^–960 cm^−1^. Below the minimum frequency of the template IR spectrum (61 cm^−1^), the spectrum was extended by a constant to 0 cm^−1^. At near infrared frequencies >3800 cm^−1^, i.e., above the maximum frequency of measurement, the spectrum was extended by a flat profile in the interband region 3800 cm^−1^–50,000 cm^−1^, and by a free electron behaviour ~ω^−4^ beyond 50,000 cm^−1^ [20, 21]. This extrapolation procedure yielded reflectance spectra R(*ω*) extended to the full frequency range. Phases *θ*(*ω*) were then retrieved from these spectra by the regularised Kramers–Kronig formula[22].

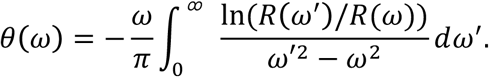

The KKT was performed by evaluating the integral with a numerical Maclaurin summation formula [21] in the range 61 cm^−1^–3800 cm^−1^, and with exact expressions[20] in the ranges 0–61 cm^−1^ and >3800 cm^−1^. The absorption coefficient *α*(*ν*) that measures the intensity absorbed in the sample per unit length is

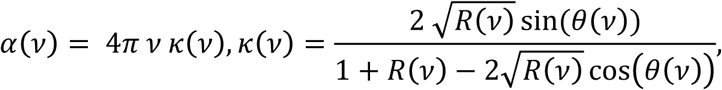

where 
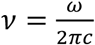
 is the frequency in cm^−1^ (inverse wavelength), and *k*(*ν*) is the extinction coefficient. For sample thicknesses *d* much larger than 
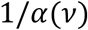
, absorbance spectra are simply proportional to the absorption coefficient: 
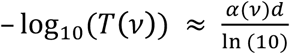
.

**Figure 3:**
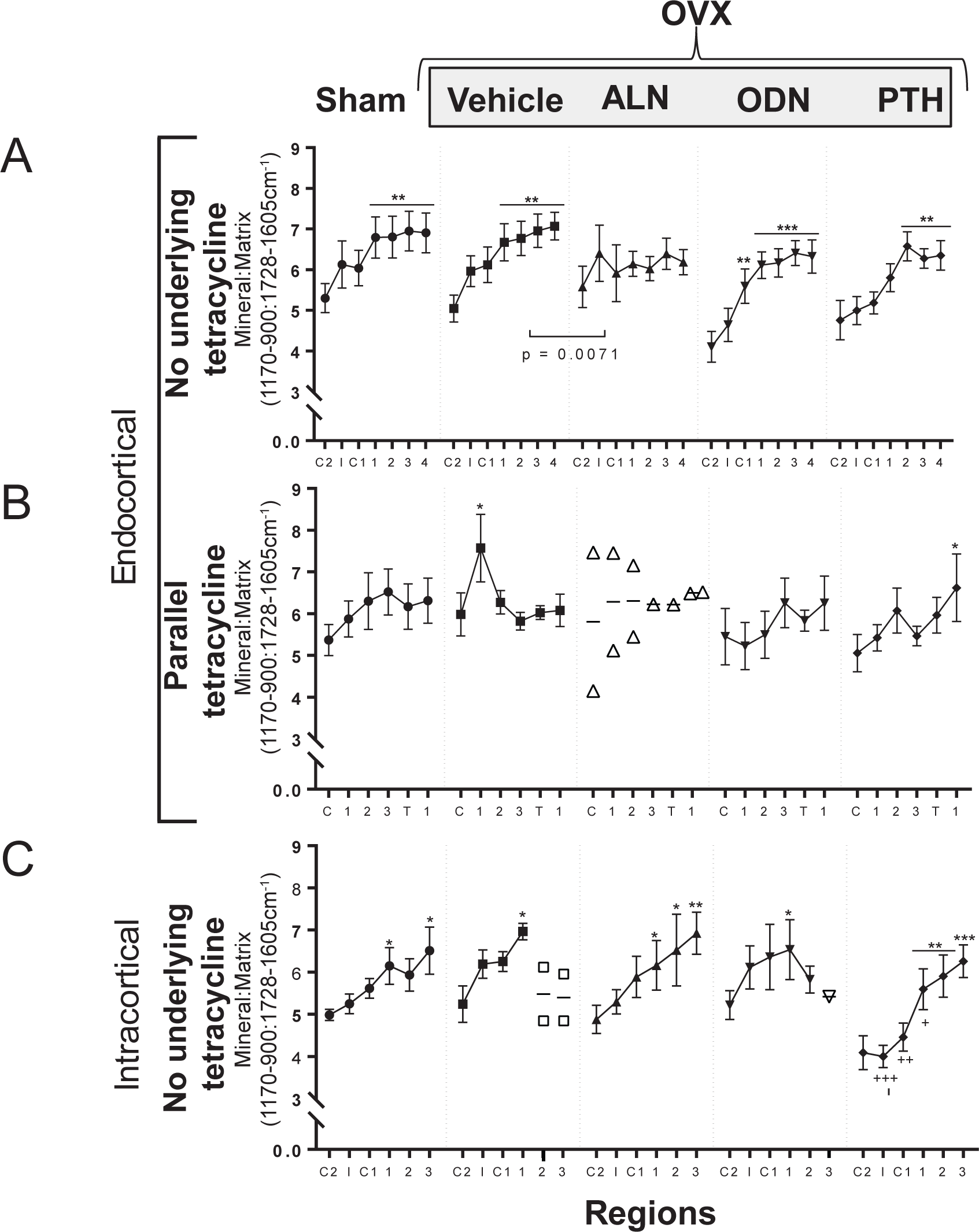
Mineral:matrix ratio in sham-operated and ovariectomized (OVX) rabbits treated with vehicle, alendronate (ALN), odanacatib (ODN) or parathyroid hormone (PTH) measured at endocortical surfaces without underlying tetracycline, and with underlying parallel tetracycline (B) and at intracortical surfaces without underlying tetracycline (C). Spectra were collected at calcein labels (C1 and C2), intermediate zone between the two calcein labels (I), tetracycline label (T) and deeper regions of bone, as defined in Figure 1. *, p<0.05, **, p<0.01, ***, p<0.001 vs C2 or C1; +p<0.05, ++p<0.01, +++p<0.001 vs same region of OVX+vehicle; p values shown with square brackets indicate significant differences in slope of the curve. Data is mean ± SEM; n= 3-9 per group; where n<3, individual data points are shown.

**Table 2:**
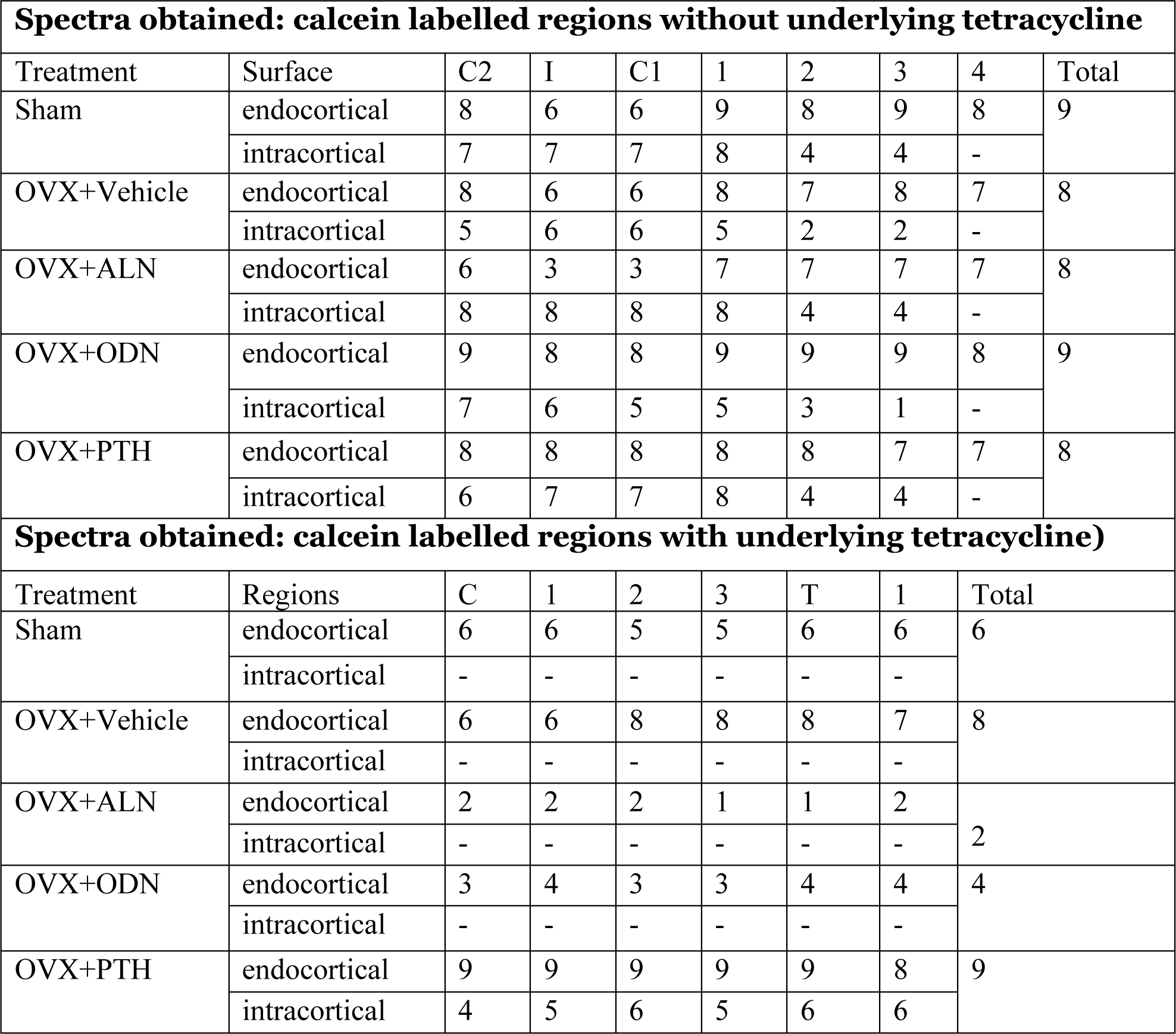
Number of spectra obtained in regions defined according to their location relative to the bone surface. See Figure 1 for additional information. C2 = calcein label closest to the bone edge (last injected label), I = intermediate region between calcein labels. C1= calcein label furthest from the bone edge. T = tetracycline label. 1-4 are regions obtained at increasing depth into the cortical bone. Total samples = the total number of specimens assessed; − = no samples exhibited this region.

| Spectra obtained: calcein labelled regions without underlying tetracycline |  |  |  |  |  |  |  |  |  |
| --- | --- | --- | --- | --- | --- | --- | --- | --- | --- |
| Treatment | Surface | C2 | I | C1 | 1 | 2 | 3 | 4 | Total |
| Sham | endocortical | 8 | 6 | 6 | 9 | 8 | 9 | 8 | 9 |
|  | intracortical | 7 | 7 | 7 | 8 | 4 | 4 | - |  |
| OVX+Vehicle | endocortical | 8 | 6 | 6 | 8 | 7 | 8 | 7 | 8 |
|  | intracortical | 5 | 6 | 6 | 5 | 2 | 2 | - |  |
| OVX+ALN | endocortical | 6 | 3 | 3 | 7 | 7 | 7 | 7 | 8 |
|  | intracortical | 8 | 8 | 8 | 8 | 4 | 4 | - |  |
| OVX+ODN | endocortical | 9 | 8 | 8 | 9 | 9 | 9 | 8 | 9 |
|  | intracortical | 7 | 6 | 5 | 5 | 3 | 1 | - |  |
| OVX+PTH | endocortical | 8 | 8 | 8 | 8 | 8 | 7 | 7 | 8 |
|  | intracortical | 6 | 7 | 7 | 8 | 4 | 4 | - |  |
| Spectra obtained: calcein labelled regions with underlying tetracycline) |  |  |  |  |  |  |  |  |  |
| Treatment | Regions | C | 1 | 2 | 3 | T | 1 | Total |  |
| Sham | endocortical | 6 | 6 | 5 | 5 | 6 | 6 | 6 |  |
|  | intracortical | - | - | - | - | - | - |  |  |
| OVX+Vehicle | endocortical | 6 | 6 | 8 | 8 | 8 | 7 | 8 |  |
|  | intracortical | - | - | - | - | - | - |  |  |
| OVX+ALN | endocortical | 2 | 2 | 2 | 1 | 1 | 2 | 2 |  |
|  | intracortical | - | - | - | - | - | - |  |  |
| OVX+ODN | endocortical | 3 | 4 | 3 | 3 | 4 | 4 | 4 |  |
|  | intracortical | - | - | - | - | - | - |  |  |
| OVX+PTH | endocortical | 9 | 9 | 9 | 9 | 9 | 8 | 9 |  |
|  | intracortical | 4 | 5 | 6 | 5 | 6 | 6 |  |  |

Absorbance spectra were analysed as previously described [16]. Briefly, spectra obtained by KKT transformation were baseline corrected using a 3-point baseline at wavenumbers 1800, 800, and the closest minimum to 1200 cm^−1^ using OPUS 7.2 as previously described. Residual MMA absorbance peaks were then subtracted using the relevant MMA reference spectrum for each sample by iterative manual subtraction using OPUS 7.2. Spectroscopic parameters calculated were integrated peak areas of the following: Phosphate (mineral) (1170 cm^−1^−900 cm^−1^), amide I (matrix) (1728 cm^−1^–1605 cm^−1^), amide II (1605 cm^−1^–1512 cm^−1^) and carbonate (890 cm^−1^–837 cm^−1^), as previously described [16, 23, 24]. Ratios of these integrated peak areas were calculated as follows: mineral:matrix ratio (1170 cm^−1^−900 cm^−1^: 1728 cm^−1^–1605 cm^−1^), carbonate:mineral ratio (890 cm^−1^–837 cm^−1^: 1170 cm^−1^−900 cm^−1^) and amide I:II ratio (1728 cm^−1^–1605 cm^−1^: 1605 cm^−1^–1512 cm^−1^). These parameters, when measured at increasing depth into maturing bone reflect bone matrix maturation, specifically: mineral accrual (mineral:matrix ratio), carbonate substitution into the hydroxyapatite matrix (carbonate:mineral) and collagen molecular compaction in its perpendicular axis (amide I:II ratio)[16]. A reduction in amide I:II ratio could also reflect collagen molecular stretching in its parallel axis, but in the context of mineralizing bone, this seems the less likely interpretation of this parameter.

### Statistical analysis

The software POWER by Lawrence Erlbaum and Associates (Hillsdale, NJ; copyright 1988) was used to calculate the sample size necessary to detect expected surgery and treatment-related differences. We specified the need to have a 90% chance of finding a significant difference.

Mechanical testing and histomorphometric data were analysed by one-way ANOVA with Fishers LSD post-hoc tests using Statview software (SAS Institute Inc., Cary, NC). FTIRM data were analysed by two-way ANOVA with Fishers LSD post-hoc tests using Prism 7 to determine the overall treatment effects, treatment effects within bone-age matched regions, and changes across regions within each treatment group. Linear regressions were performed to test whether slopes were significantly different in treatment groups compared to vehicle-treated OVX samples. P<0.05 was considered significant.

## Results

### In-Life Study

At the start of the study, the mean body weights of the Sham and OVX rabbits 6 months post-OVX were significantly different (p<0.01): Sham (3.62 kg ± 0.06) and OVX (3.78 kg ± 0.06). Rabbits were maintained on a restricted diet to combat body weight changes between Sham and OVX-rabbits. Rabbits given PTH at 15 μg/kg/5x per week exhibited a group mean body weight loss of 0.18 gm, representing a 4.7% loss in body weight during the experiment, but there was no loss in body weight in the ALN or ODN treated groups. There were no significant serum calcium concentration differences among groups.

### Treatment effects on bone strength and bone formation on endocortical, intracortical and periosteal surfaces

After 10 months of treatment, *ex vivo* assessment (not corrected for body weight) indicated greater aBMD in ODN and ALN-treated rabbits, but not PTH-treated rabbits (Figure 2A), consistent with previous reports [25]. Biomechanical testing showed significantly greater toughness and work-to-failure in PTH and ODN treated femora compared to vehicle treated OVX controls (**Table 1**); ALN did not significantly modify mechanical strength. No other mechanical testing parameters were modified by treatment (**Table 1**).

As expected, OVX induced an increase in periosteal mineralising surface (PsMS/BS); this was suppressed by ALN, and further promoted by PTH treatment (Figure 2B). Although PTH and ODN have both been reported to promote periosteal bone formation in other species[7, 26], in this study there was insufficient double labelled surface to measure periosteal mineral appositional rate (PsMAR) in any treatment group; of all samples collected, only 2 PTH-treated samples had sufficient double label to measure PsMAR (Figure 2B).

ALN treatment suppressed endocortical bone formation (Figure 2C,D). Most ALN samples exhibit insufficient double label to allow endocortical mineral appositional rate (EcMAR) measurement (10/14 specimens, see **Table 1**), as expected given the known effects of ALN on suppressing bone formation. This meant that, when prepared for FTIRM analysis, most ALN-treated samples exhibited only single calcein labels on the endocortical surface (**Table 2**) and very few samples showed any surfaces where calcein was detected on surfaces containing underlying tetracycline label (modelling bone). In contrast, PTH treatment increased endocortical (Ec) MS/BS and Ec.MAR, and consequently increased bone formation rate at this region (Figure 2C-E). ODN treatment neither suppressed nor stimulated bone formation on the periosteal or endocortical surfaces (Figure 2C,D).

PTH-treated rabbits demonstrated more intracortical Haversian bone formation than all other groups of rabbits (**Table 1**). This included a Haversian mineralising surface (H.MS/BS) of approximately 10%, and Haversian mineral appositional rate (H.MAR) and Haversian BFR/BS (H.BFR/BS) equivalent to endocortical bone of Sham-operated and OVX rabbits. These parameters could not be measured consistently in any other group because of insufficient intracortical double label. Only the PTH-treated group showed any calcein labelling on intracortical bone with underlying tetracycline label (modelling surfaces).

### Synchrotron-based FTIRM defines the normal pattern of bone mineral accrual in Sham-operated and OVX rabbits

Synchrotron-based FTIRM analysis of recently formed endocortical bone in Sham-operated rabbits demonstrated the same three characteristics of bone matrix maturation that we previously observed in recently formed bone on the murine periosteum [16]: (1) mineral accrual, (2) carbonate substitution, and (3) collagen compaction, indicated by amide I:II suppression (Figure 3A, 4A, 5A). Carbonate substitution, indicated by carbonate:mineral ratio, gradually increased with increasing bone age at all sites measured (Figure 4A-C). Other parameters showed site-specific differences. In Sham animals, the increase in mineral:matrix ratio with increasing depth into the bone (Figure 3A) was similar in both intracortical and endocortical sites of short-term bone formation (bone forming surfaces lacking underlying tetracycline label (F < 4mo)) (Figure 4C). In contrast, in long-term bone forming sites (F > 4mo), where double calcein labels were incorporated above a parallel tetracycline label) there was no significant increase in mineral:matrix ratio (Figure 3B). Although lowering of amide I:II with increasing bone age was significant in both endocortical regions, it was not significant in intracortical bone (Figure 5A-C).

**Figure 4:**
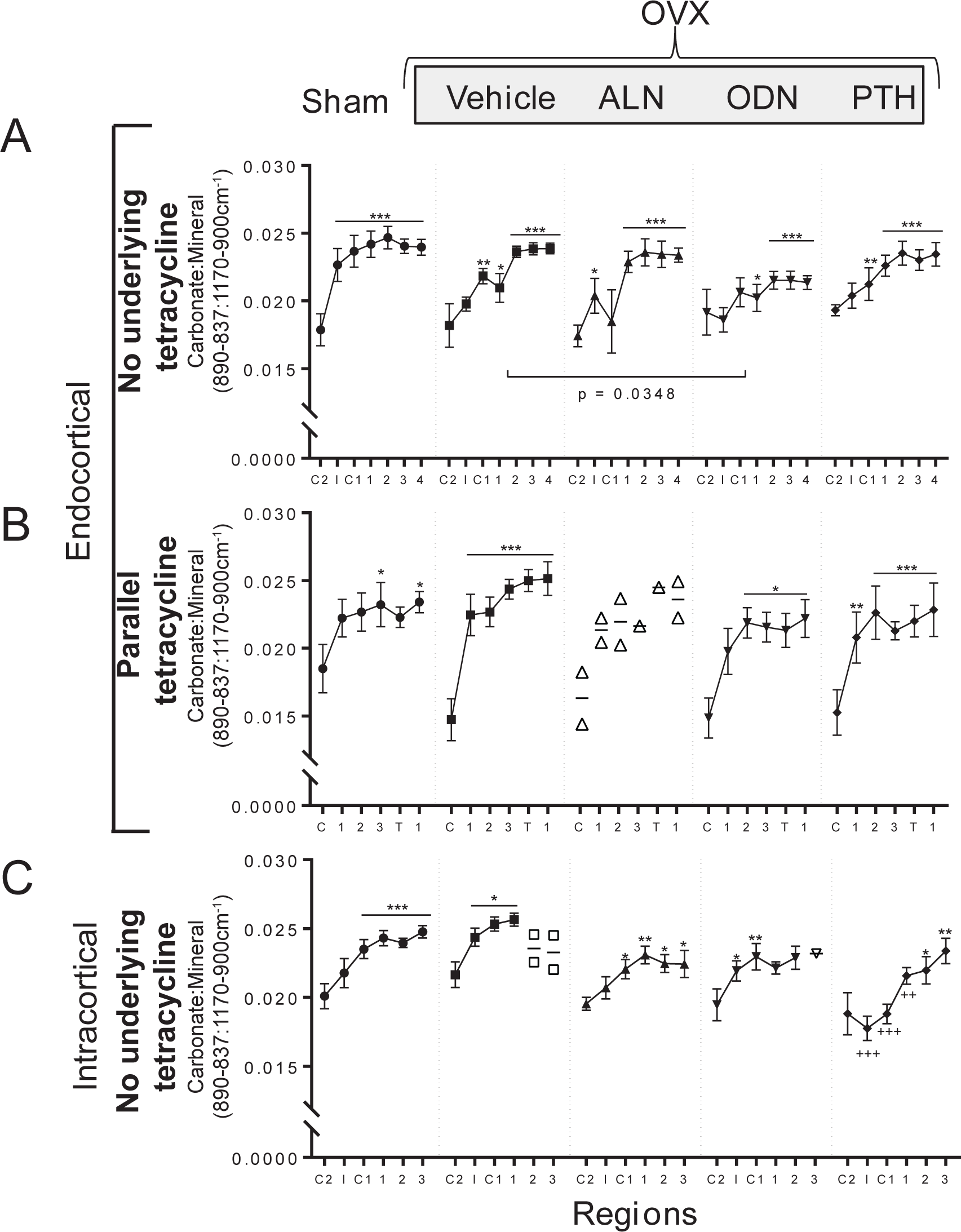
Carbonate:mineral ratio in sham-operated and ovariectomized (OVX) rabbits treated with vehicle, alendronate (ALN), odanacatib (ODN) or parathyroid hormone (PTH) measured at endocortical surfaces without underlying tetracycline (F <4mo, A), and with underlying parallel tetracycline (B) and at intracortical surfaces without underlying tetracycline (C). Spectra were collected at calcein labels (C1 and C2), intermediate zone between the two calcein labels (I), tetracycline label (T) and deeper regions of bone, as defined in Figure 1. *, p<0.05, **, p<0.01, ***, p<0.001 vs C2 or C1; +p<0.05, ++p<0.01, +++p<0.001 vs same region of OVX+vehicle; p values shown with square brackets indicate significant differences in slope of the curve. Data is mean ± SEM; n= 3-9 per group; where n<3, individual data points are shown.

**Figure 5:**
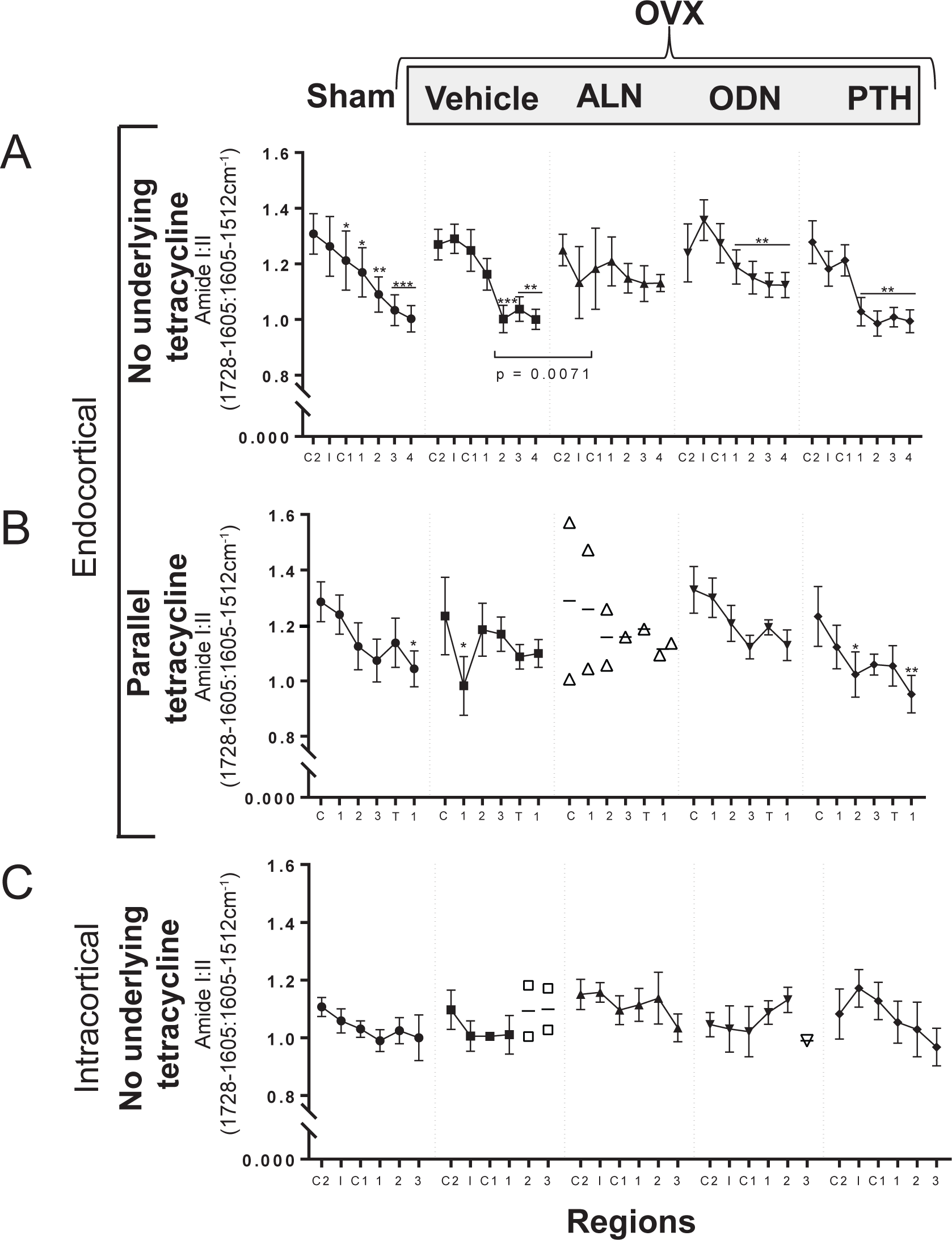
Amide I:II ratio in sham-operated and ovariectomized (OVX) rabbits treated with vehicle, alendronate (ALN), odanacatib (ODN) or parathyroid hormone (PTH) measured at endocortical surfaces without underlying tetracycline (A), and with underlying parallel tetracycline (B) and at intracortical surfaces without underlying tetracycline (C). Spectra were collected at calcein labels (C1 and C2), intermediate zone between the two calcein labels (I), tetracycline label (T) and deeper regions of bone, as defined in Figure 1. *, p<0.05, **, p<0.01, ***, p<0.001 vs C2 or C1; p values shown with square brackets indicate significant differences in slope of the curve. Data is mean ± SEM; n= 3-9 per group; where n<3, individual data points are shown.

In OVX rabbits, the increase in mineral:matrix and carbonate:mineral ratios, and the reduction in amide I:II ratio in endocortical bone with short-term bone formation was very similar to Sham rabbits (Figure 3A, 4A, 5A). As observed in Sham rabbits, OVX samples did not show a significant increase in mineral:matrix ratio in endocortical bone with long-term bone formation (Figure 3A), although carbonate:mineral significantly increased (Figure 4A). In intracortical bone, initial accrual of mineral and carbonate was observed, but it was not possible to obtain sufficient spectra for analysis of deep regions of bone matrix, due to the high porosity of the cortical bone in OVX samples, so a comparison of the deep regions could not be made in this region.

### ALN treatment disrupts matrix maturation in endocortical, but not intracortical bone

In endocortical bone, the pattern of bone matrix maturation was changed by ALN treatment (Figure 3A). Due to suppression of bone formation by ALN, sufficient spectra could only be obtained in endocortical bone without underlying tetracycline label (F > 4mo, Figure 3A). In this region, although the normal pattern of carbonate substitution was maintained (Figure 4A), neither the increase in mineral:matrix ratio, nor the decrease in amide I:II ratio observed in Sham and OVX samples were detected in ALN-treated samples (Figure 3A, 5A); the slopes of these lines were significantly lower than in OVX samples measured in the same region (both p=0.007). In contrast, ALN-treated intracortical bone showed a continual increase in mineral:matrix and carbonate:mineral ratios, as observed in Sham animals (Figure 3C,4C).

### ODN treatment suppressed mineral:matrix accrual and modified matrix carbonate substitution

ODN treatment was associated with two changes in matrix maturation in intracortical bone and in endocortical bone lacking underlying tetracycline (for clarity direct comparisons with OVX are shown in Figure 6). These changes were: (1) a lower level of carbonate substitution in both regions (Figure 6B,D), and (2) lower mean mineral:matrix ratio in the endocortical region (Figure 6A). ODN-treated rabbits had significantly lower mineral:matrix and carbonate:mineral ratios (Figure 6A,B), and higher amide I:II ratio (Figure 6C) than vehicle-treated OVX rabbits in endocortical regions without underlying tetracycline; the latter two differences were most distinct in deeper regions of bone, suggesting carbonate substitution and collagen molecular compaction were impaired in the late stages of bone matrix maturation in the presence of ODN. In the intracortical region, there were no significant differences in mineral:matrix or amide I:II ratios (Figure 6D,F), but carbonate:mineral levels were significantly lower, both overall, and in bone regions closest to the osteonal centre (Figure 6E).

**Figure 6:**
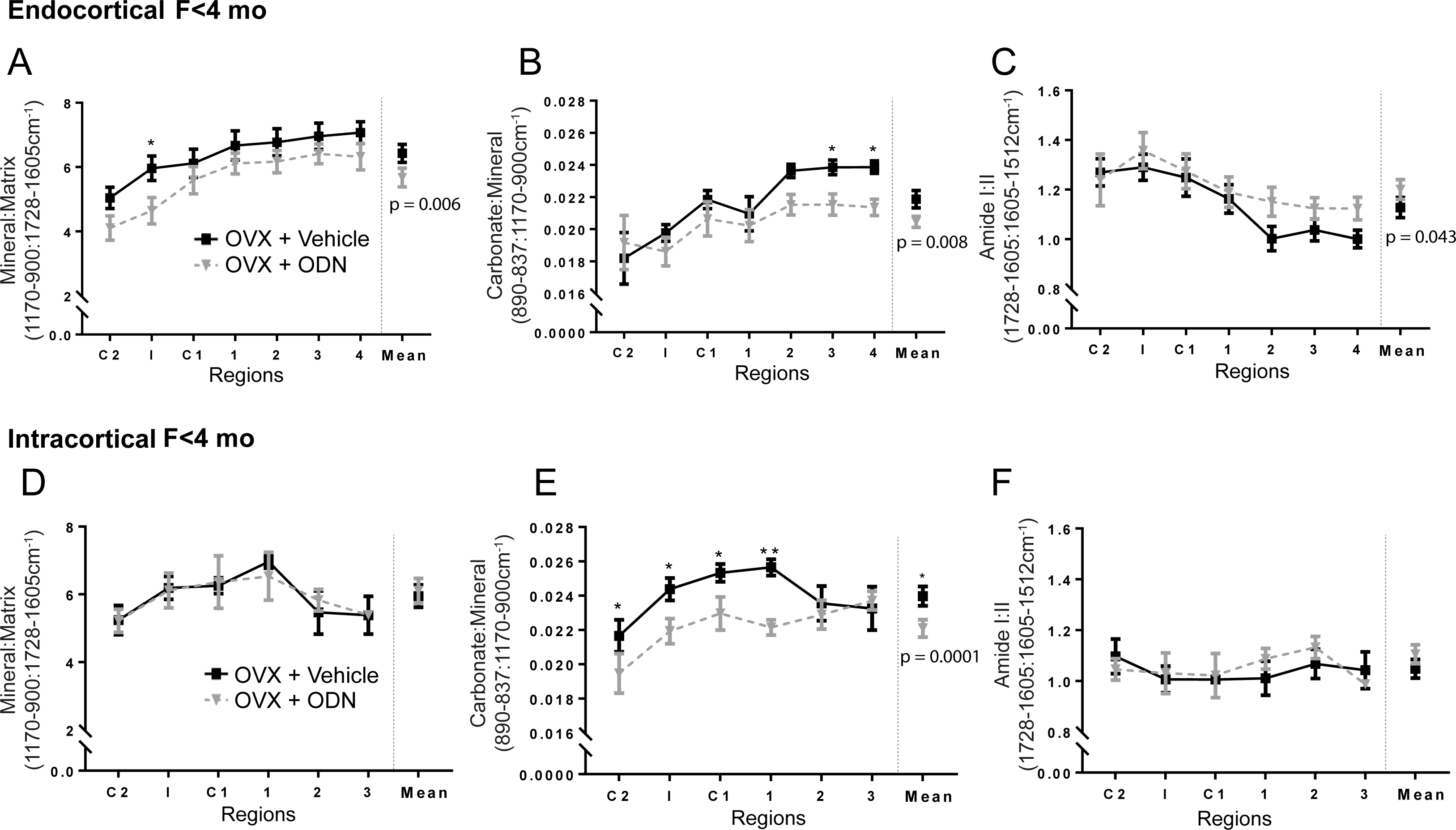
Direct comparison of the changes in mineral:matrix, carbonate:mineral and amide I:II ratios with increasing depth into the bone matrix in ODN and vehicle-treated OVX rabbits on endocortical (A-C) and intracortical (D-F) surfaces without underlying tetracycline. This is the same data as shown in Figures 4-6, but on the same axes. *, p<0.05, **, p<0.01 vs OVX+vehicle in the same region. Data is mean ± SEM; n= 8-9 per group. At the right-hand side of each graph is the mean value for each group and the overall p value for treatment effect from a two-way ANOVA analysis.

### Bone formed during PTH treatment shows normal bone maturation dynamics, except in intracortical bone

PTH treatment did not significantly modify the normal progression of bone maturation in endocortical regions compared to OVX-vehicle or Sham samples (Figures 3,4,5). In contrast, in intracortical bone, although PTH-treated bone reached the same mineral:matrix level in deepest regions of bone, it had significantly lower mineral:matrix and carbonate:mineral ratios than vehicle-treated OVX samples in newly formed bone nearest the calcein labels (Figure 3C, 4C).

Although bone formation with underlying parallel tetracycline label (modelling-based bone formation) was rarely observed in intracortical bone in Sham, OVX, ALN-treated, and ODN-treated samples (Table 2), PTH treatment induced bone formation on these surfaces, indicating modeling-based bone formation in intracortical bone. We therefore also assessed mineral maturation in that region. This newly formed bone showed the same pattern of mineral accrual (increased mineral:matrix ratio), carbonate substitution (increased carbonate:mineral) and reduced amide I:II ratio with increasing bone age (Figure 7) as Sham samples. The slope elevations for all these parameters were significantly less than those of the same parameters on endocortical modelling surfaces (mineral:matrix ratio, p<0.001; carbonate:mineral ratio, p=0.0175; amide I:II ratio, p=0.0028). This suggests that new bone formed in response to PTH treatment in intracortical spaces may mature less rapidly than elsewhere.

**Figure 7:**
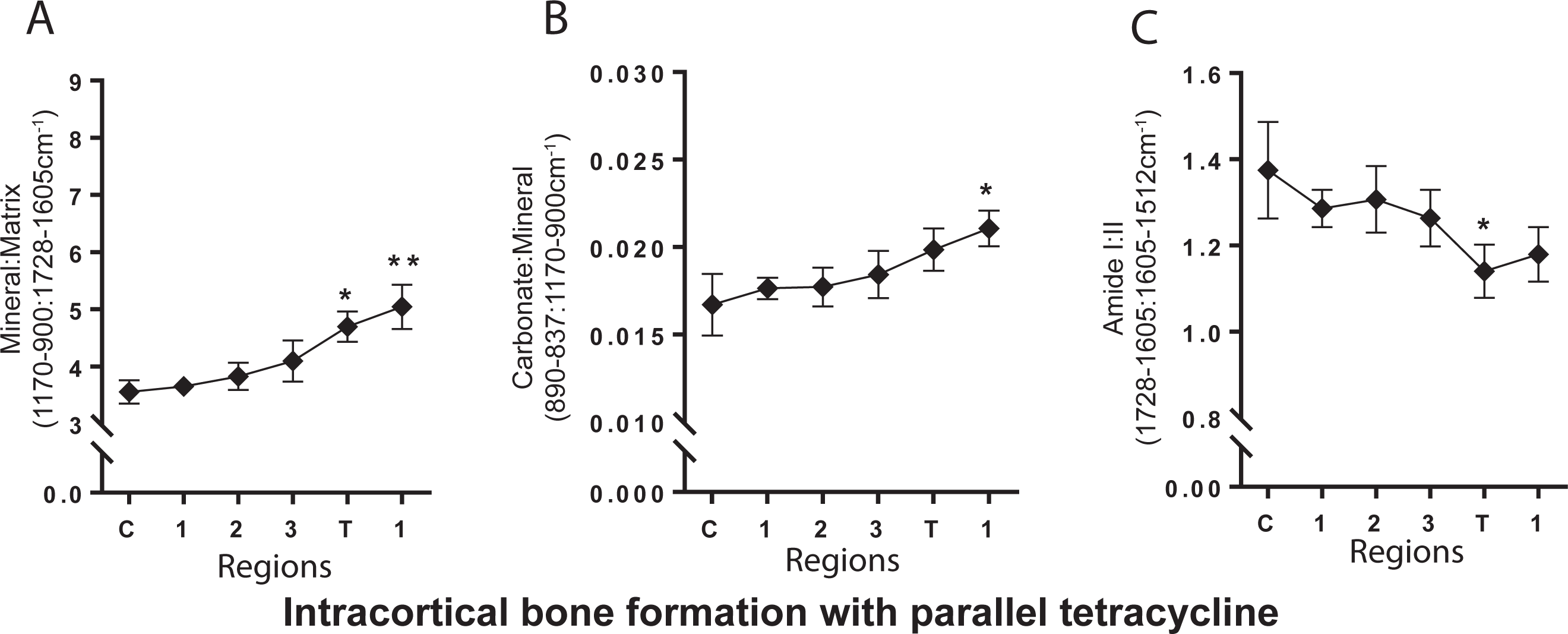
Mineral:matrix (A), carbonate:mineral (B) and amide I:II (C) ratios on intracortical surfaces at sites of bone formation with underlying tetracycline label in OVX rabbits treated with PTH. *, p<0.05, **, p<0.01 vs region C. Data is mean ± SEM; n= 6 per group.

## Discussion

This study provides three main insights into the way bone formation and bone matrix maturation are modified by ODN and PTH treatment: (1) PTH treatment stimulates bone formation not only on endocortical bone surfaces, but also promotes bone formation in intracortical bone. (2) Mineral maturation in PTH-treated rabbits resulted in mature bone with normal composition in all regions, but in intracortical bone, the initial phase of mineralization was slower than in controls. (3) The processes of mineral accrual and carbonate substitution were slower in ODN-treated bone. It also shows that ALN treatment may dissociate carbonate substitution from other aspects of bone matrix maturation, although this data was limited by the low level of bone formation induced by ALN.

This is the first report showing that PTH treatment stimulates bone modelling in intracortical bone. Bone formation in Haversian osteonal bone with underlying parallel tetracycline label (modelling-based bone formation) was only observed in PTH-treated bones. The finding that PTH stimulates bone formation on these surfaces is consistent with previous reports that PTH stimulates bone formation on both quiescent (modelling) and remodelling trabecular and endocortical bone [13, 27-29], and that PTH may reactivate bone lining cells to promote bone formation [30] without requiring additional osteoclast-derived coupling factors to promote bone formation[31]. Our finding of intracortical bone formation even on modelling surfaces suggests that PTH can fill, or partially fill, intracortical pores without requiring prior resorption. This suggests that new bone formation can occur in intracortical pores in which resorption is not stimulated, and contrasts with the increased intracortical porosity reported in both animal and human studies [15, 32, 33]. This provides new information about how PTH strengthens bone, and may explain why weekly (rather than daily) PTH administration did not increase cortical porosity in non-OVX rabbits [34].

PTH treatment did not alter mineral accrual, carbonate substitution or amide I:II ratio in endocortical bone, consistent with our previous observations in murine periosteum[16]. However, in intracortical regions, PTH-treated bone accrued mineral less rapidly. This was true both when modelling-based intracortical bone formation was compared with either modelling-based endocortical bone formation, or with intracortical bone that lacked underlying tetracycline label. The latter site would represent a mixture of remodelling-based bone formation (tetracycline has been resorbed), and recently initiated modelling-based formation (on a quiescent, non-labelled surface). The low mineral:matrix level in newly-formed PTH-treated bone indicates that, at least initially, mineral accrual is slower at this site during PTH treatment. The reason for slower initial mineral accrual is not clear; it may relate to a limited capacity of the mineral reservoir that is unable to fully mineralise the larger than usual volume of newly deposited osteoid. The slow mineral accrual is temporary; mineral:matrix levels continued to increase in deeper regions of bone (see Figure 3C, PTH); this indicates that PTH treatment has restored the normal bone maturation pattern that was lost in OVX rabbits (see Figure 3C, vehicle).

This increase in bone formation in intracortical osteons in PTH-treated samples provides an observation that contrasts to the increased intracortical resorption and porosity reported in PTH-treated rabbits [34], and monkeys [15, 33]. It should be noted that the PTH-induced increase in cortical porosity is not uniform, and occurs preferentially near the endocortical surface, in a way that limits negative biomechanical effects [35]. There is clearly variation in responses of intracortical pores to PTH, which may depend on mechanical loading, or on the stage of basic multicellular unit (BMU)-based remodelling existing at each site when PTH is supplied. For example, at sites where osteoclasts are already present, increased resorption would occur, and at sites where bone formation is already occurring (such as those with underlying tetracycline label), modelling-based bone formation would occur.

ODN treatment resulted in lower overall mineral content and carbonate substitution in endocortical bone, and a significantly higher amide I:II ratio in endocortical regions without underlying tetracycline. ODN also suppressed the rate of mineral accrual: mineral:matrix ratio was lower in newly formed (calcein labelled) ODN-treated bone. A low level of carbonate substitution was also observed, both in endocortical and intracortical bone. In intracortical bone, this occurred without any change in either mineral:matrix or amide I:II, suggesting that ODN-treatment changes the normal relationship between these bone matrix components. Previous studies of bone composition with ODN treatment focused on calcium content by back-scattered electron imaging (qBEI) in OVX monkeys treated with ODN [36], and observed no changes. This suggests that while ODN does not alter calcium levels, it does change the nature of the hydroxyapatite crystal, by modifying the proportion of calcium to phosphate and the level of carbonate substitution within the hydroxyapatite lattice. While it is not known how this occurs, it may be caused by ODN’s action to reduce collagen removal at the resorption surface prior to laying down new bone; this may explain why it was only observed in remodelling, rather than modelling, bone. Alternately, since cathepsin K is expressed by osteocytes [37], it may relate to unknown cathepsin K-mediated actions in osteocytes to modify hydroxyapatite composition in the surrounding new bone matrix. High mineral:matrix and carbonate:phosphate ratios are associated with greater bone hardness and indentation modulus at the nanomechanical level [38]. In contrast, at the whole bone level, higher carbonate substitution has been reported in fractured femurs [39]. The ultimate effect of reduced carbonate substitution during cathepsin K inhibition is not known, but since ODN treatment increased bone toughness, we suggest that the reduced carbonate substitution may be beneficial.

Cathepsin K inhibition by ODN has been reported to suppress bone resorption without blocking bone formation [7-9], a finding partially upheld by the present study, where histomorphometry showed no significant reduction in EcMAR or PsMAR. Previous work in OVX monkeys has reported increased bone formation on both periosteal and endocortical surfaces in the central femur [40], but we did not see this, suggesting a species difference in the effect of ODN on periosteal bone formation. We also observed less fluorochrome labelling in intracortical bone in ODN-treated rabbits compared to endocortical bone, which is consistent with previous work in monkeys showing that ODN suppressed intracortical bone formation[8, 41].

ALN suppresses osteoclast-mediated bone resorption and reduces bone turnover rate [42], an observation we confirmed on both intracortical and endocortical surfaces. It has been reported that ALN decreases mineral heterogeneity in trans-iliac bone biopsies [43-46] and increases the mean degree of mineralisation, quantified by back-scatter electron microscopy, due to prolonged secondary mineralisation [47]. An FTIRM study using long term treatment and multiple fluorochrome labels observed no alteration in the rate at which newly formed trabecular bone becomes mineralised in the presence of ALN [4]; our findings in intracortical bone in the present study is consistent with that earlier finding. In intracortical bone, we also detected that carbonate substitution and the reduction in amide I:II ratio associated with bone matrix aging occurred at a normal rate. We did not observe this on endocortical surfaces. In those regions ALN-treatment blunted both the increase in mineral:matrix ratio and the decrease in amide I:II ratio. In contrast, we reliably detected an increase in carbonate:mineral with bone age on the same spectra from ALN-treated samples. This suggests carbonate substitution may occur independently of mineral accrual and collagen molecular compaction, at least in ALN-treated bone.

In conclusion, maturation of cortical bone matrix is altered by ODN, PTH and ALN treatment. We confirmed the inhibitory action of ALN on bone formation and showed that it may also dissociate carbonate substitution from other aspects of bone matrix maturation. ODN treatment slowed mineral accrual, and limited carbonate substitution; if alternative therapeutic methods of inhibiting cathepsin K activity are developed, this may have significant effects on skeletal composition in the long term. Finally, PTH treatment stimulated bone formation in intracortical bone, even on modeling surfaces, and while the accrual of mineral proceeded normally in PTH-treated endocortical bone, it was slower in newly formed intracortical bone.

## Acknowledgements

The authors gratefully acknowledge technical assistance from Christine Jun, extensive advice and consultation on the KKT method from Alvin Acerbo and Lisa Miller at Brookhaven National Laboratories, and very helpful critique of the manuscript from T. John Martin. This work was funded by an investigator-initiated grant from MSD to NAS. NAS is funded by an NHMRC Senior Research Fellowship. sFTIRM was undertaken on the Infrared Microspectroscopy beamline at the Australian Synchrotron, part of ANSTO. St. Vincent’s Institute receives funds from the Victorian Government’s Operational Infrastructure Support Program.

Study design: NAS and LTD. Study conduct: NAS, CV, PB, TAP, BLP, KRB. Data collection: CV, TAP, BLP. Data analysis: CV, TAP, BLP, NAS. Data interpretation: CV, PB, KRB, LTD, NAS. Drafting manuscript: CV, PB and NAS. Revising manuscript content: all authors. Approving final version of manuscript: All authors. NAS takes responsibility for the integrity of the data analysis.

